# White matter alterations at pubertal onset

**DOI:** 10.1101/128165

**Authors:** Sila Genc, Marc L Seal, Thijs Dhollander, Charles B Malpas, Philip Hazell, Timothy J Silk

## Abstract

Recent neurodevelopmental research supports the contribution of pubertal stage to local and global grey and white matter remodelling. Little is known, however, about white matter microstructural alterations at pubertal onset. This study investigated differences in white matter properties between pre-pubertal and pubertal children using whole brain fixel-based analysis (FBA) of the microscopic density and macroscopic cross-section of fibre bundles.

Diffusion-weighted imaging data were acquired for 74 typically developing children (*M*=10.4, *SD*=0.43 years, 31 female) at 3.0T (60 diffusion gradient directions, b-value=2800 s/mm^2^). Group comparisons of fibre density (FD) and fibre cross-section (FC) were made between age-matched pre-pubertal and pubertal groups, and post-hoc analyses were performed on regions of interest (ROIs) defined in the splenium, body and genu of the corpus callosum.

Significant fixel-wise differences in FD were observed between the pubertal groups, where the pubertal group had significantly higher FD compared with age-matched pre-pubertal children, localised to the posterior corpus callosum. Post-hoc analyses on mean FD in the corpus callosum ROIs revealed group differences between the pubertal groups in the splenium, but not body or genu.

The observed higher apparent fibre density in the splenium suggests that pubertal onset coincides with white matter differences explained by increasing axon diameter. This may be an important effect to account for over pubertal development, particularly for group studies where age-matched clinical and typical populations may be at various stages of puberty.

**Abbreviations:**

BMI: Body mass index
CFE: Connectivity-based fixel enhancement
DTI: Diffusion tensor imaging
DWI: Diffusion-weighted imaging
FA: Fractional anisotropy
FBA: Fixel-based analysis
FC: Fibre cross-section
FD: Fibre density
FDC: Fibre density and cross-section
FOD: Fibre orientation distribution
FWE: Family-wise error
GLM: General linear model
HPA: hypothalamic-pituitary-adrenal
HPG: Hypothalamic-pituitary-gonadal
JHU: John’s Hopkins University
MRI: Magnetic resonance imaging
PDS: Pubertal development scale
SEIFA: Socioeconomic indexes for areas
SES: Socio-economic status
TE: Echo-time
TR: Repetition time
WASI: Wechsler abbreviated scale of intelligence

## 1. Introduction

Puberty is a critical period of development, marking the transition from childhood to reproductive maturity (Dorn et al., 2006). Adrenarche describes the earliest phase of puberty, beginning between the ages of 6 – 9 in females, and approximately 1 year later in males (Grumbach and Styne, 1998). It is the period in which physical changes begin to manifest. Endocrine events tied to puberty such as activation of the hypothalamic-pituitary-adrenal (HPA) and -gonadal (HPG) axes leads to increased production of adrenal and gonadal steroid hormones, which have been shown to enter the blood brain barrier and initiate remodelling of the brain in animal models (Sisk and Foster, 2004).

Quantifying these hormone levels in puberty, however, is inherently limited by individual variability and diurnal pattern of hormone concentrations, making it difficult to infer pubertal stage (Ankarberg and Norjavaara, 1999). The Pubertal Development Scale (PDS) is a non-invasive self or parent-report measure of pubertal stage that is highly correlated with hormones and physical exam (Shirtcliff et al., 2009).

A growing body of evidence suggests that the timing of pubertal onset can lead to differential patterns of grey and white matter development evidenced by Magnetic Resonance Imaging (MRI) (Blakemore et al., 2010; Byrne et al., 2016; Ladouceur et al., 2012). In the context of grey matter, there is converging evidence of a positive relationship between pubertal stage and pituitary volume (Whittle et al., 2012; Wong et al., 2014), global grey matter density (Peper et al., 2009), and amygdala volume (Goddings et al., 2014; Neufang et al., 2009).

In addition to physical indices of pubertal maturation, endogenous adrenal and sex steroid hormone levels have been associated with grey and white matter structure. In early puberty, corpus callosum body volume is higher with increased luteinizing hormone (LH) levels (Peper et al., 2008). Testosterone levels are associated with increased white matter volume (Perrin et al., 2008), as well as sex-dependent relationships in grey matter surface area (Herting et al., 2015) and corticospinal structure (Pangelinan et al., 2016).

Diffusion MRI techniques have allowed the in-vivo quantification of properties of white matter microstructure during this dynamic period of brain development. A number of diffusion tensor imaging (DTI) studies have revealed greater microstructural organisation with pubertal advancement (Asato et al., 2010; Herting et al., 2012; Menzies et al., 2015). Whilst DTI is the most commonly used technique to assess white matter microstructure, metrics such as fractional anisotropy (FA) are non-specific at distinguishing between white matter properties such as axon density, glial infiltration, crossing fibres, and partial voluming. These are all separate physio-anatomical properties of white matter that must be disentangled to fully understand the nature of white matter changes across development (Beaulieu, 2009).

Since chronological age and pubertal stage are highly correlated, highly sensitive as well as specific models that interrogate white matter microstructure at the fibre level are necessary to disentangle ‘pubertal age’, or pubertal contributions to white matter microstructure, from chronological age. Despite these advancements, disentangling age-related mechanisms of brain development from pubertal processes remains a challenge.

Recent advances have introduced more specific measures of axon density and fibre bundle cross-section (Groeschel et al., 2016). Neurite density index from NODDI (Zhang et al., 2012) can characterise the density of neurites by restricted diffusion by modelling the intra-cellular space (Sepehrband et al., 2015). Whilst more sensitive than DTI metrics to age-related differences in the developing brain (Genc et al., 2017), there has been some debate regarding its interpretation in areas in multiple crossing fibres, and the underlying assumptions of the model (Lampinen et al., 2017; Novikov, 2016).

Fixel-based analysis (FBA) (Raffelt et al., 2012; Raffelt et al., 2017) is a recently developed diffusion MRI analysis technique that uses fibre-population specific information to estimate fibre density and morphology for individual white matter fibre populations. A *fixel* is defined as a *fibre*-population within a *voxel*, and as such FBA allows for whole-brain comparisons of fibre-specific white matter properties. It produces metrics that assess fibre density (FD), fibre cross-section (FC), and the combined effect of fibre density and cross-section (FDC), where FD for a given fibre population is proportional to the volume of the intra-axonal compartment. FC reflects the cross-sectional area of the white matter fibre bundle, and FDC reflects the combined effect of FD and FC. Increases in FD can represent an increase in axonal count, or denser packing of axons, in a given fibre area, whereas increases in FC can suggest greater cross-sectional area of a fibre bundle.

In this study, we implement fixel-based analysis to estimate and compare white matter fibre density and morphology across two age-matched groups of children: pre-pubertal (no evidence of maturation of secondary sexual characteristics), and pubertal (evidence of pubertal maturation).

## 2. Methods

### Participants

This paper reports on a subsample of typically developing children recruited as part of the Neuroimaging of the Children’s Attention Project study (see Silk et al. (2016) for a detailed protocol). Briefly, children were recruited from 43 socio-economically diverse primary schools distributed across the Melbourne metropolitan area, Victoria, Australia. MR imaging was performed on 86 typically developing children. Of those, 8 participants were excluded owing to missing or motion-affected diffusion MRI data, and a further 4 participants were excluded owing to missing pubertal data, resulting in a total of 74 children aged 9.5 – 11.9 years being included in the current study. This study was approved by The Royal Children’s Hospital Melbourne Human Research Ethics Committee (HREC #34071)

As part of the assessment procedure, a number of participant characteristics were determined based on child and parent report. Socio-economic status (SES) was determined using the Socio-Economic Indexes for Areas (SEIFA). General intellectual ability was estimated using the Wechsler Abbreviated Scale of Intelligence (WASI) matrix reasoning sub-test (Wechsler, 1999). Height and weight were measured using the average of two consecutive measurements to calculate a Body-Mass index (BMI) (kg/m^2^). Participant characteristics are summarised in *Table 1*.

**Table 1:**
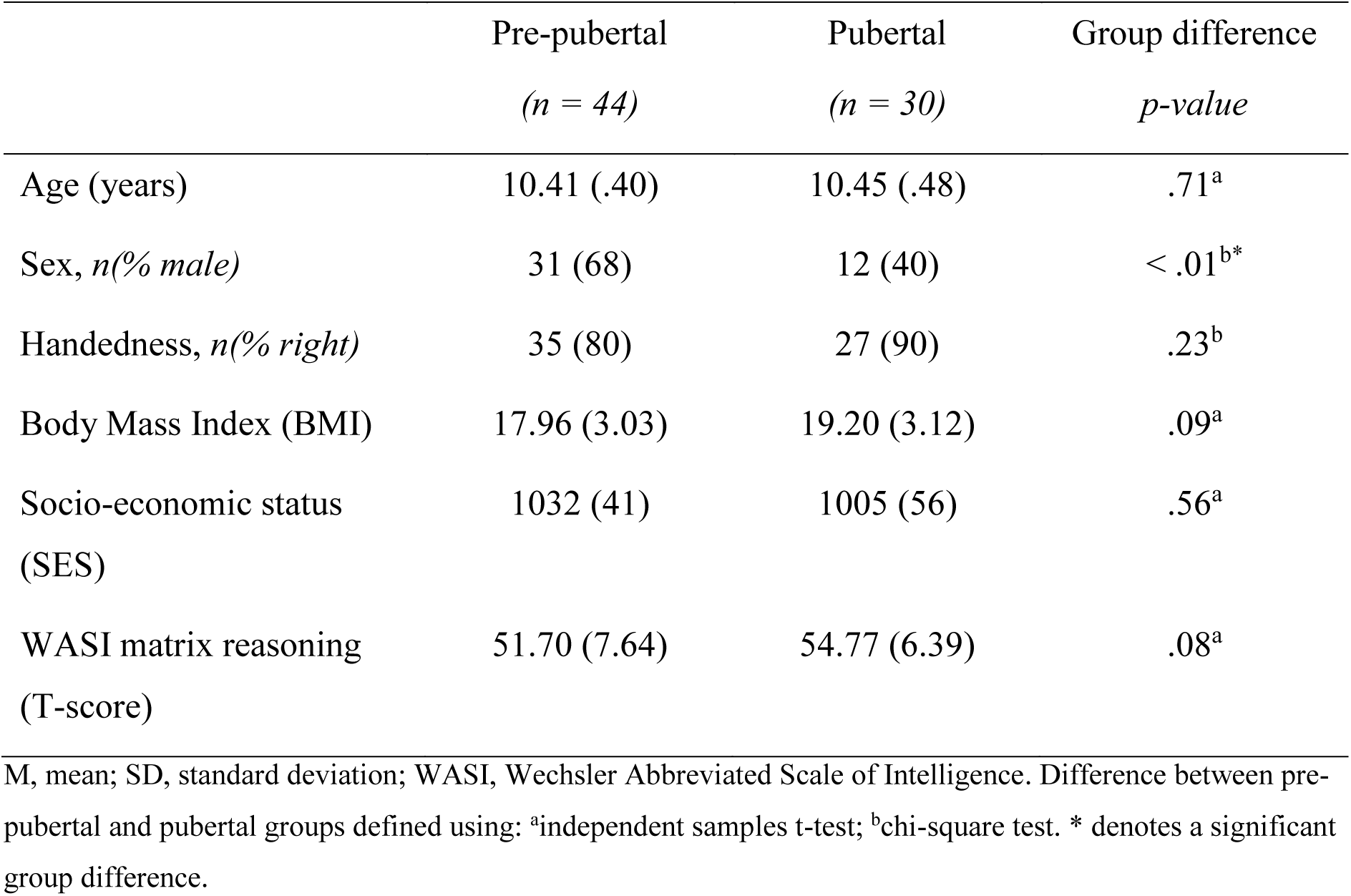
Demographics and participant characteristics by pubertal group. Values reported as *M* (*SD*) unless otherwise stated.

### Pubertal classification

Pubertal development was assessed using the Pubertal Development Scale (PDS) (Petersen et al., 1988). The PDS is a measure that assesses pubertal stage and is highly correlated with hormonal measures and physical exam by a trained physician, rendering it a reliable measure of pubertal maturation (Shirtcliff et al., 2009). For the current study, the primary caregiver was asked to rate their child’s physical development on a four-point scale. This included questions assessing the presence of characteristics phenotypical of pubertal onset such as deepening of voice and presence of facial hair in boys, and breast development and menarche for females. A combined PDS-Shirtcliff (PDSS) score was calculated (Shirtcliff et al., 2009) by taking the mean of the adrenarche and gonadarche scores generated from the syntax described in the aforementioned study. The PDSS scores in our sample ranged from 1 – 3.5, where children with a score of 1 had no physical signs of pubertal onset, and children with a score of 1.5 - 3.5 had some phenotypic characteristics of pubertal onset. Subsequently, the sample was divided into two groups: *pre-pubertal* (children with PDSS = 1) and *early-pubertal* (children with PDSS ≥ 1.5). For ease of explanation, the early-pubertal group (PDSS ≥ 1.5) will be referred to as the *pubertal* group.

### Image acquisition and processing

Diffusion-weighted imaging (DWI) data were acquired on a research-dedicated 3T Siemens Tim Trio MRI scanner (Erlangen, Germany) at The Melbourne Children’s campus, Melbourne, Australia. Data were acquired using 60 diffusion gradient directions, b-value = 2800 s/mm^2^, 2.4 × 2.4 × 2.4 mm^3^ isotropic voxel size, Echo-time/Repetition time (TE/TR) = 110/3200 ms, and 4 volumes without diffusion-weighting (b = 0). Diffusion data were pre-processed using the MRtrix3 (version 0.3.15,https://github.com/MRtrix3) software package which included: denoising (Veraart et al., 2016); motion, eddy current and susceptibility distortion correction (Andersson and Sotiropoulos, 2016); bias field correction (Tustison et al., 2010); and global (group-wise) intensity normalisation.

### Fixel-based analysis

Fixel-based analysis (FBA) was performed in accordance with a recommended pipeline (Raffelt et al., 2017). Firstly, data were spatially up-sampled by a factor of two, and fibre-orientation distributions (FODs) were estimated for each participant using a group average response function. An unbiased study-specific FOD template was created using 28 participants with equal numbers of subjects from each pubertal group, matched for sex.

Whole brain tractography was performed on the FOD template where 20 million tracts were generated, and spherical-deconvolution informed filtering of tractograms (SIFT) was implemented to filter this down to 2 million tracts, the density of which corresponds to the fibre density present in the data (Smith et al., 2013). Subsequently, FBA output metrics FD, FC and FDC were calculated.

An FOD-based direction encoded colour (DEC) map (Dhollander T, 2015), was used as a reference backdrop to inspect the fibre orientation of individual fixels. Colouring is according to tract orientation (left-right: red, inferior-superior: blue, anterior-posterior: green).

### Statistical analyses

General Linear Models (GLMs) were computed for permutation-based testing of FBA metrics, covarying for age and sex. We compared FD, FC and FDC between the groups using *Connectivity-based fixel enhancement* (CFE) in MRtrix (Raffelt et al., 2015). The FBA results reported here were generated using 5,000 permutations and reached family-wise error (FWE) corrected statistical significance at *p_FWE_* < .05. Main effect of sex, and age, on FBA metrics was examined, with PDS group included in the model. The interaction between sex, and age, with FBA metrics was also assessed, with PDS group included in the model in order to establish homogeneity of regression. Mean values were extracted from the significant group-difference cluster and subjected to post-hoc categorical regression onto PDSS score using *CATreg* in the SPSS software (IBM Corporation, version 23).

Secondary to the whole-brain analysis, we investigated a potential gradient of white matter development along the corpus callosum by generating regions of interest (splenium, body, and genu of corpus callosum) for further analysis. First, the JHU-ICBM FA template (Smith et al., 2004) was registered to the FOD template, and the transformation was applied to the three corpus callosum masks from the JHU white matter labels atlas. Secondly, these masks were overlaid onto the FOD template with fixel overlay, and each mask was manually edited for concordance with the template (to correct for subtle misregistrations), as well as ensuring only single-fibre (i.e. single fixel) voxels were included. Mean FD was calculated across each region for all participants, and a region by group interaction was also investigated using a repeated measures MANOVA. Finally, mean values were compared between groups for each region. Standardised effect sizes are reported as Hedge’s g (Hedges, 1981).

## 3. Results

### Participant characteristics

The pre-pubertal and pubertal group did not differ in age, handedness, BMI, SES, or WASI matrix reasoning T-score (*Table 1*). Sex distribution significantly differed between the groups, as there was a higher proportion of girls in the pubertal group compared to the prepubertal group.

### Fixel-based analysis

Fixel-based analyses revealed higher FD in the pubertal group, compared with the prepubertal group (*p_FWE_* < .001). The region of significantly higher fibre density was localised to the splenium of the corpus callosum, and this group difference is visualised in *Figure 1*, thresholded at *p_FWE_* < .05. No regions had significantly lower fibre density in the pubertal (non-significant negative contrast). There were no significant group differences in FC or FDC.

**Figure 1:**
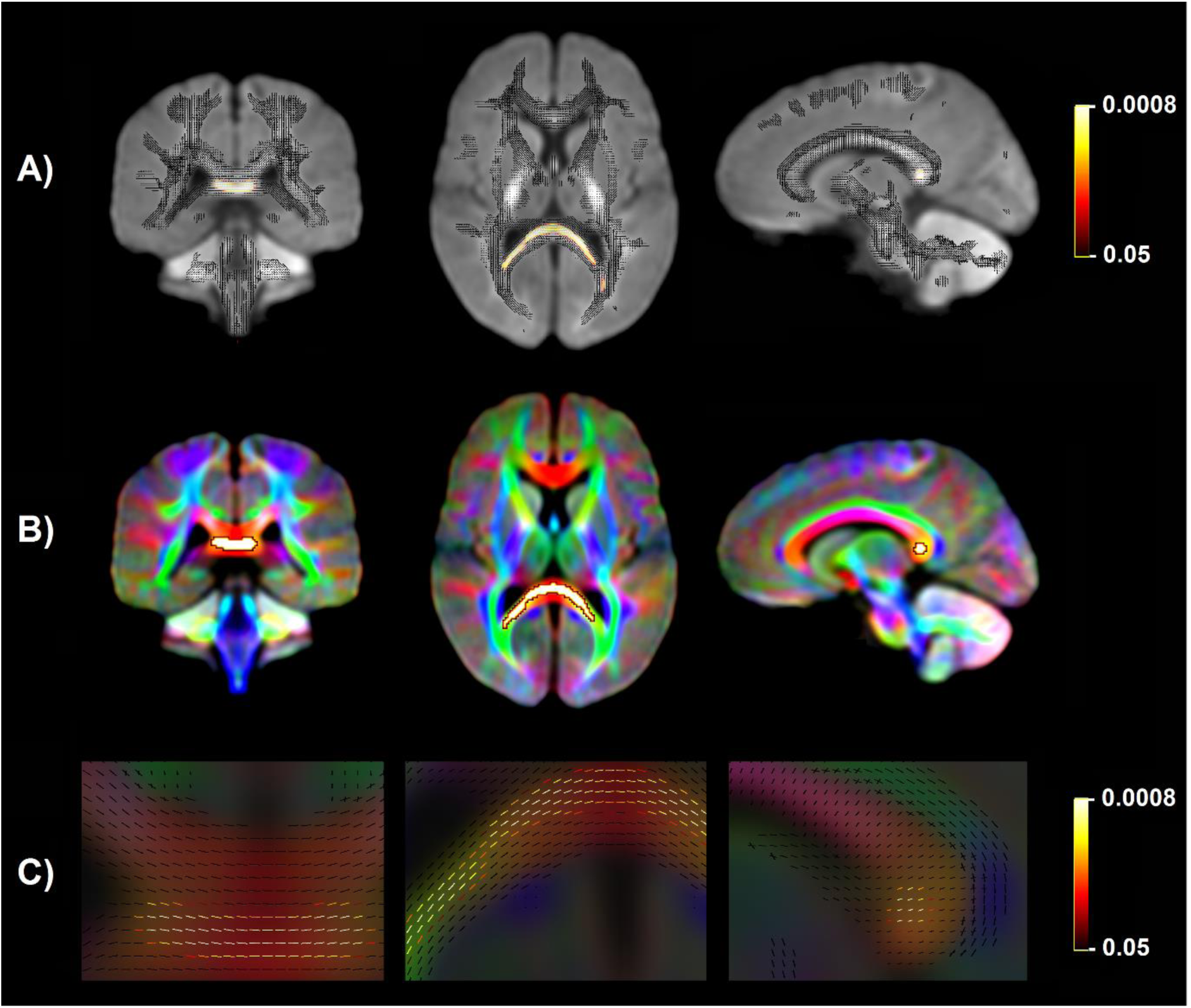
Fixels with significantly higher fibre density (FD) in the pubertal group compared with the pre-pubertal group. A) Fixel analysis mask (black), with colour-coded significant fixels (*p_FWE_* < .05) overlaid on the FOD template. B) Significant fixels converted to a voxel-map (white) overlaid on the DEC map. C) Magnification of colour-coded significant fixels to view fibre direction in the significant region, overlaid on the DEC map. Colour bars in A) and C) represent p-value thresholds, where most significant fixels (*p_FWE_* < .001) are coloured in yellow.

Sex by group, and age by group interactions were non-significant across FD, FC and FDC (*p_FWE_* > .05). Therefore homogeneity of regression was not violated for the FBA, and use of sex and age as covariates in our GLM tests were valid assumptions.

There was no main effect of age on FBA metrics across the groups, and only a main effect of sex on FC was found, however given group differences were only present in FD this result was not further explored.

Post-hoc analyses on mean FD in the significant ROI revealed group differences between the pubertal groups. Mean FD in the pre-pubertal group (*M* = .897, *SD* = .061) was lower than in the pubertal group (*M* = .971, *SD* = .069). This difference was statistically significant, using age and sex as covariates (*t* = 4.28, *df* = 70, *p* < .0001) (*Figure 2a*). These group differences persisted when splitting the groups up by sex and comparing FD between pre-pubertal and pubertal children in males and females separately, covarying for age (*Figure S1*). Pubertal males had higher FD than pre-pubertal males (*t* = 3.41, *df* = 40, *p* = .0015), and pubertal females had higher FD than pre-pubertal females (t = 2.66, *df* = 28 *p* = .0127), which is consistent with the overall findings.

**Figure 2:**
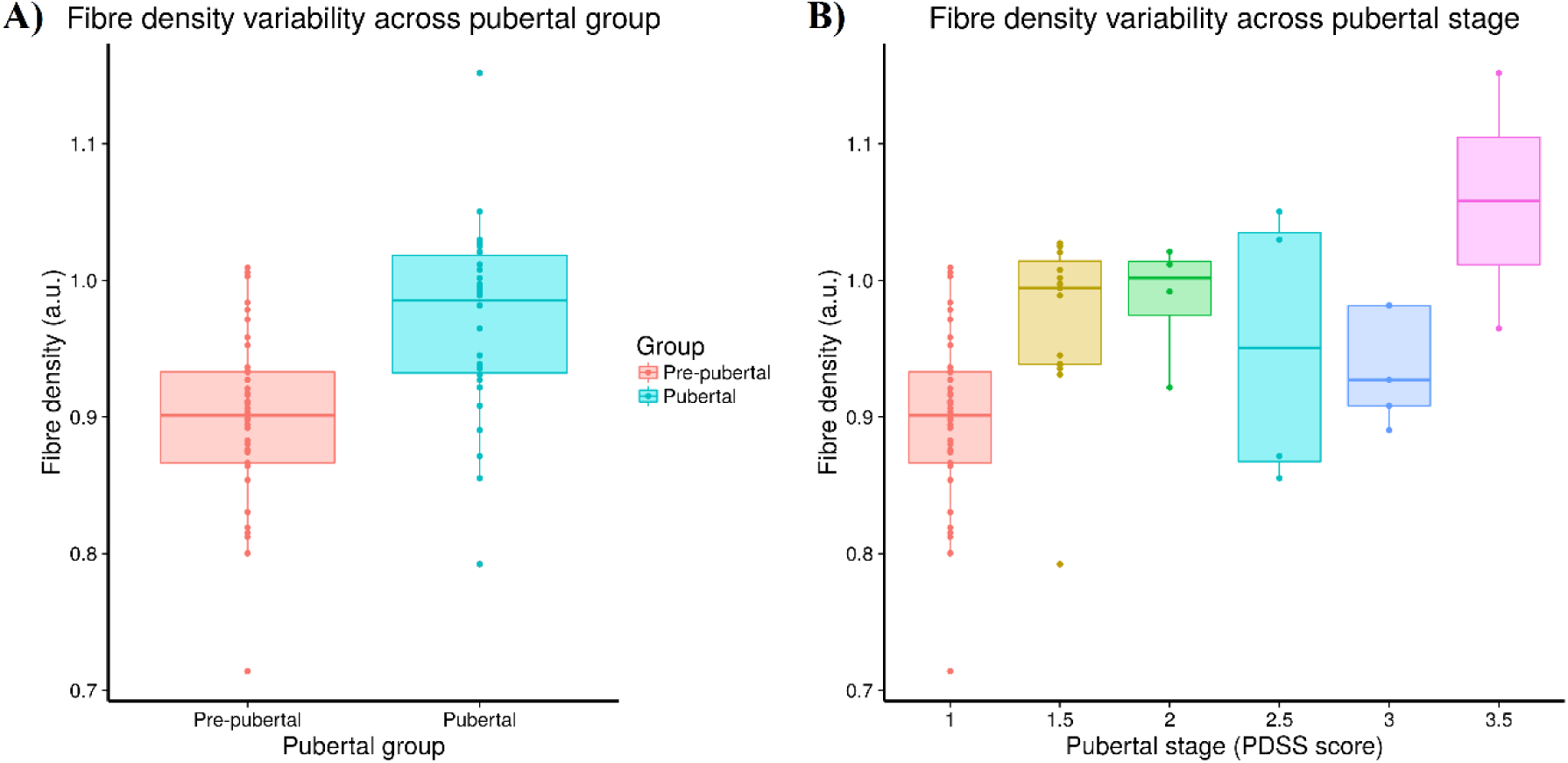
Spread of fibre density across pubertal stage: A) Spread of FD in significant region by pubertal group; B) Spread of FD in significant region by PDSS score. FD values are represented in arbitrary units (a.u.).

As the number of participants in PDSS stage 2 or above was small (*Figure 2b*), we performed exploratory categorical regressions to investigate the relationship between PDSS score and FD, with age as a covariate. Across the entire sample, FD was a significant predictor of PDSS score (β = .46, *p* = .005). Further categorical regression was performed in each sex separately, and FD was a significant predictor of PDSS scores in males (β = .47, *p* = .005) and females (β = .50, *p* = .002) separately. The direction of all relationships were consistent with the FBA findings, with higher FD associated with greater PDSS score.

### Corpus callosum parcellation

To examine the gradient of fibre density across the corpus callosum, we parcellated single-fibre regions of interest (ROIs) in the splenium, body and genu (*Figure 3*), as explained in the Methods section. The region by group interaction was statistically significant, Pillai’s Trace = .13, F(2, 70) = 5.212, *p* = .008, (partial η^2^ = .13).

**Figure 3:**
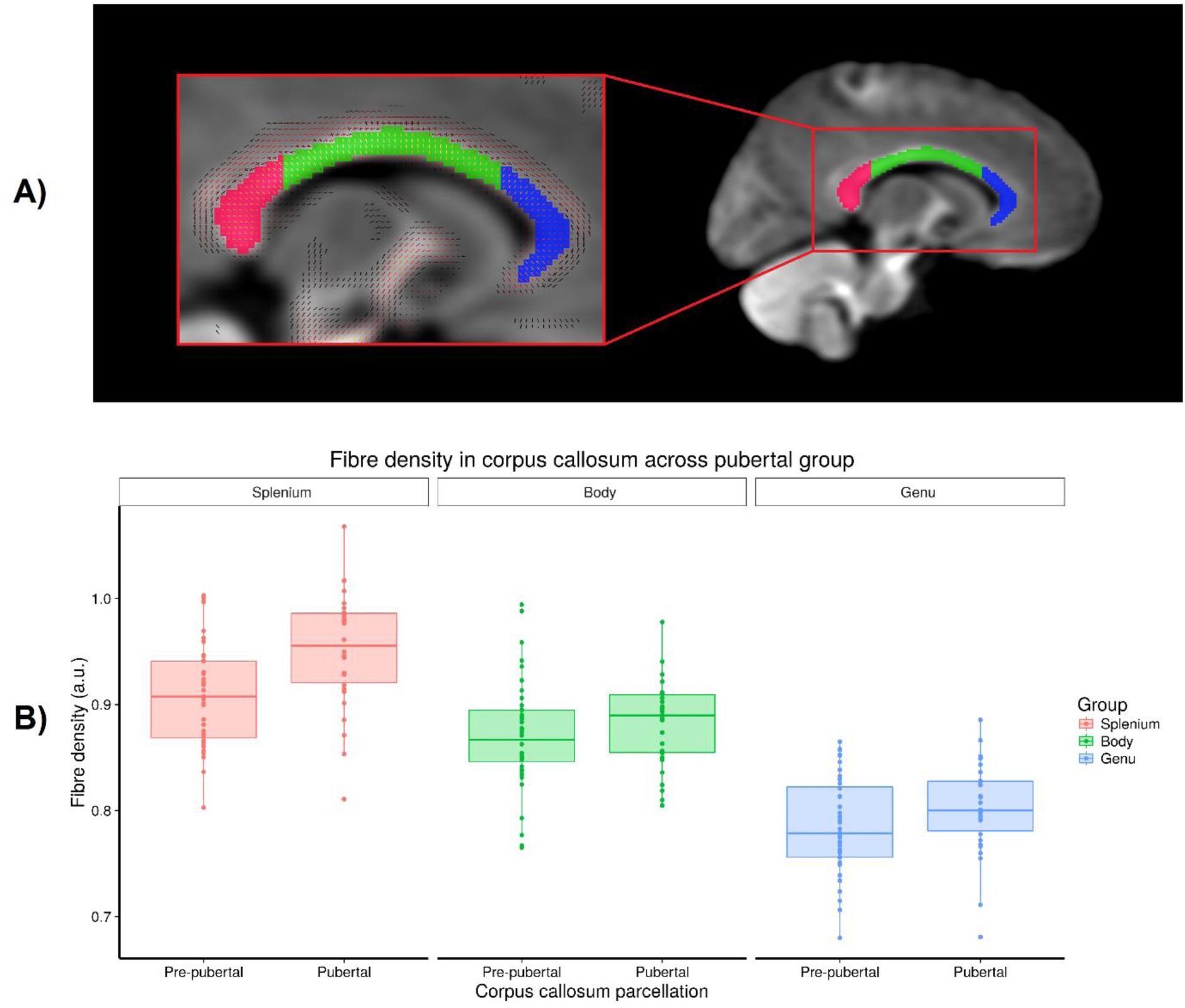
The corpus callosum parcellation where colours represent the splenium (red), body (green), and genu (blue). A) A representative slice of the corpus callosum parcellation overlaid on the FOD template, alongside magnified image with fixel overlay; B) Spread of FD in each corpus callosum section stratified by pubertal group. FD values are represented in arbitrary units (a.u.).

As shown in *Table 2*, significant differences were found for the splenium, with a large effect size. Significant differences, however, did not emerge for the body and genu.

**Table 2:**
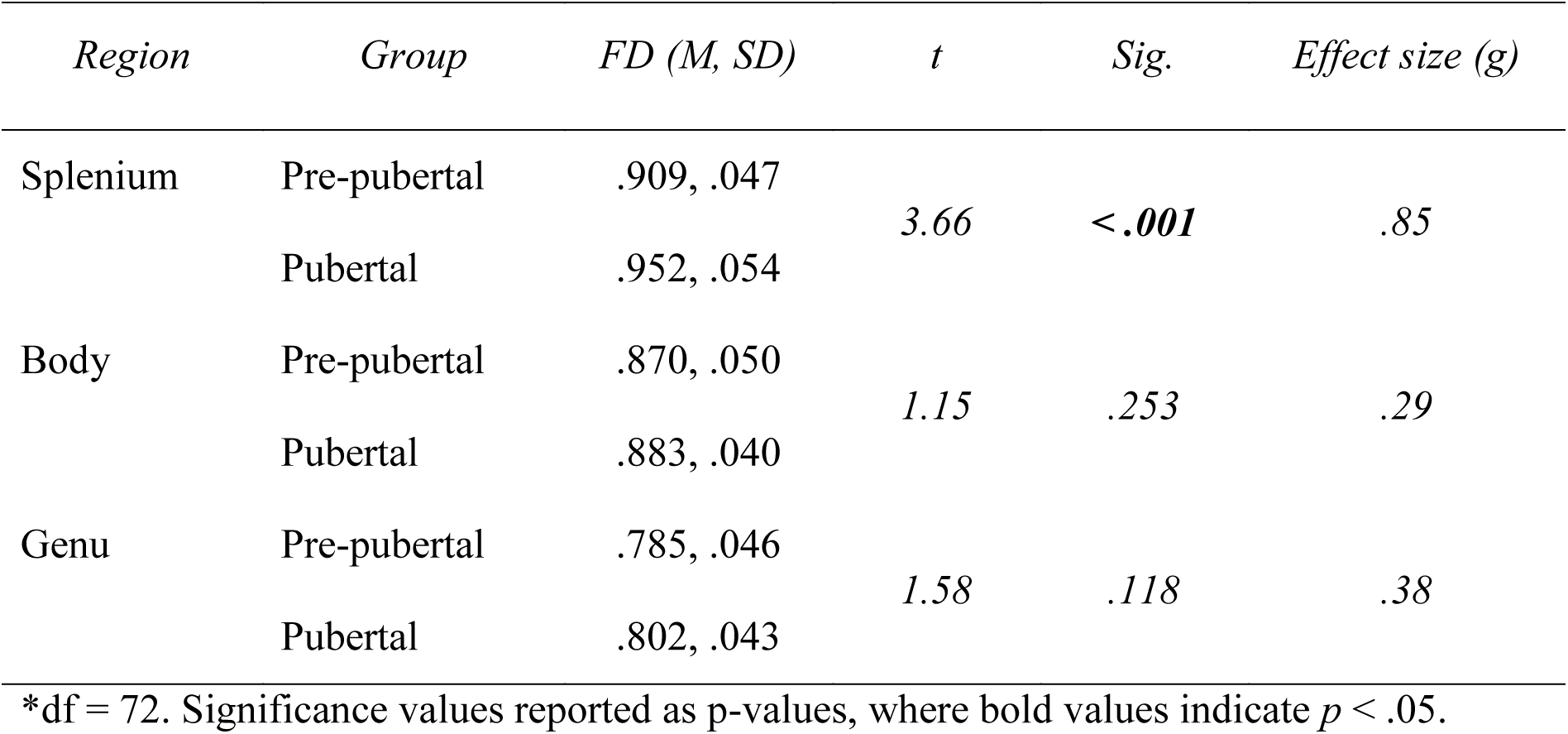
Summary of the FD comparisons between pubertal groups in the corpus callosum parcellation.

| *Region* | *Group* | *FD (M, SD)* | *t* | *Sig.* | *Effect size (g)* |
| --- | --- | --- | --- | --- | --- |
| Splenium | Pre-pubertal | .909, .047 | 3.66 | < **.001** | .85 |
|  | Pubertal | .952, .054 |  |  |  |
| Body | Pre-pubertal | .870, .050 | 1.15 | .253 | .29 |
|  | Pubertal | .883, .040 |  |  |  |
| Genu | Pre-pubertal | .785, .046 | 1.58 | .118 | .38 |
|  | Pubertal | .802, .043 |  |  |  |

\*df = 72. Significance values reported as p-values, where bold values indicate $p < .05$ .

## 4. Discussion

This study presents the first application of the recently developed fixel-based analysis technique to investigate white matter fibre architectural differences between age-matched children at pre- and early-stages of pubertal maturation. Previous research suggests that the onset of puberty contributes to the remodelling of white matter during the transition between childhood and adolescence, and the present study shows that fibre density is specifically altered.

The fixel-wise comparison between the groups revealed higher FD in the pubertal group, compared with the pre-pubertal group, in the splenium of the corpus callosum. The group differences in FD together with the absence of group differences in FC could suggest either: (1) axons are more densely packed (i.e., increased local axon count in the bundle), or (2) individual axons have increased in diameter in a bundle of overall constant diameter, which would also result in increased total intra-axonal volume. Regardless of the specific mechanism, these findings may suggest a strengthening of the fibre bundle’s capacities and improved information transfer across the splenium.

Maturation of corpus callosum connections leads to more efficient interhemispheric communication, which is an important step in pubertal onset as a number of processes accompany pubertal progression, such as the development of executive function and emotion regulation. Age-related development of the corpus callosum has been well documented, namely volumetric expansion of the splenium (Brouwer et al., 2012), increased thickness of the splenium (Westerhausen et al., 2016), and early development of the corpus callosum indexed by FA (Lebel et al., 2012; Lebel et al., 2008). Previous studies have observed that white matter follows a posterior-to-anterior gradient of development, evidenced by histological (Yakovlev and Lecours, 1967), and DTI (Asato et al., 2010; Krogsrud et al., 2016) studies, where frontal white matter is thought to be the last to mature and reorganise in adolescence. The development of this neural circuitry relates to a number of processes that emerge in later adolescence, such as risk taking, emotion regulation and inhibitory control (Blakemore, 2012; Kwon et al., 2002).

Early evidence has suggested that the number of axons in the corpus callosum does not change over the lifespan, reaching adult levels in the early post-natal period (LaMantia and Rakic, 1990; Rakic and Yakovlev, 1968). Perrin et al. (2008) first proposed that increases in axon diameter may drive white matter changes across pubertal advancement. Indeed, certain adrenal hormones have neurobiological actions, such as dehydroepiandrosterone (DHEA) which stimulates neurite growth and neurogenesis (Maninger et al., 2009). With this converging evidence at hand, it is clear that a complex interplay of adrenal and steroid hormones act to modify white matter organisation in the brain, likely driven by increasing axon size with pubertal development.

In order to further understand callosal development in this group of children, we investigated FD in the splenium, body, and genu of the corpus callosum, and how this differs with pubertal stage. Although we found no significant pubertal group differences in the body or genu of the corpus callosum, the pattern of higher fibre density in the pubertal group across all parcellations is consistent with the contribution of pubertal onset to white matter remodelling. Previous work has linked further pubertal advancement with the corpus callosum midbody shape (Chavarria et al., 2014), and animal studies have revealed that the splenium of the corpus callosum undergoes major remodelling over puberty (Kim and Juraska, 1997), which is in line with the posterior-anterior gradient of development. Combining previous findings in the context of the present study leads us to postulate that these group differences will persist anteriorly with later stages of pubertal maturation. Longitudinal studies are required to see if these gradients are present on an individual level across specific pubertal maturation trajectories.

A strength of the current study is the relatively narrow age range of the participants. They are clustered at the cusp of pubertal onset, a time when brain remodelling is thought to begin (Byrne et al., 2016; Klauser et al., 2015; Peper et al., 2008). A limitation of other studies splitting groups into early and late stages of pubertal development is the disparity in age range between groups. Whilst useful for looking at group differences due to puberty, covarying for age may simply not be enough to disentangle age-related contributions to neurodevelopment.

The implications of this study are not solely tied to understanding trajectories in typical brain development. In neurodevelopmental and psychiatric conditions with adolescent age of onset, pubertal timing may play an important role in brain structure and organisation, so it raises the question of whether we should account for pubertal timing when comparing clinical with neurotypical populations.

### Limitations & future directions

Although the tight age range is a strength of this study to investigate early remodelling of white matter, looking into later stages of puberty is warranted to see if these differences persist and whether it follows a posterior-anterior gradient of white matter development. One limitation is the imbalance of males and females across the two groups, which is somewhat expected as girls enter puberty on average 1 year earlier than boys, so in an aged matched sample it is likely that an imbalance will be present. Future investigations could match groups based on pubertal stage as well as age, which would require larger sample sizes. Future studies should also investigate the longitudinal course of white matter change, and incorporate other biological markers of pubertal maturation such as endogenous circulating hormone levels. Longitudinal multi-modal imaging data and hormone measures are currently being collected for this cohort of children (Silk et al., 2016).

## 5. Conclusion

In conclusion, we found that white matter apparent fibre density in the splenium is associated with pubertal onset. These findings suggest that pubertal onset is associated with increasing axon diameter, compared with age-matched pre-pubertal children. Together, we present evidence that white matter development and maturation may be influenced by pubertal onset itself, rather than solely the developmental processes associated with chronological age.

## 6. Acknowledgements

This study was funded by the National Medical Health and Research Council of Australia (NHMRC; project grant #1065895). This research was conducted within the Developmental Imaging research group, Murdoch Childrens Research Institute, and the Children’s MRI Centre, The Royal Children’s Hospital, Melbourne, Victoria. It was supported by the Murdoch Childrens Research Institute, The Royal Children’s Hospital, The Department of Paediatrics at The University of Melbourne and the Victorian Government’s Operational Infrastructure Support Program. SG is supported by the Australian Government Research Training Program Scholarship.

## 10. Supplementary

**Supplementary Figure 1:**
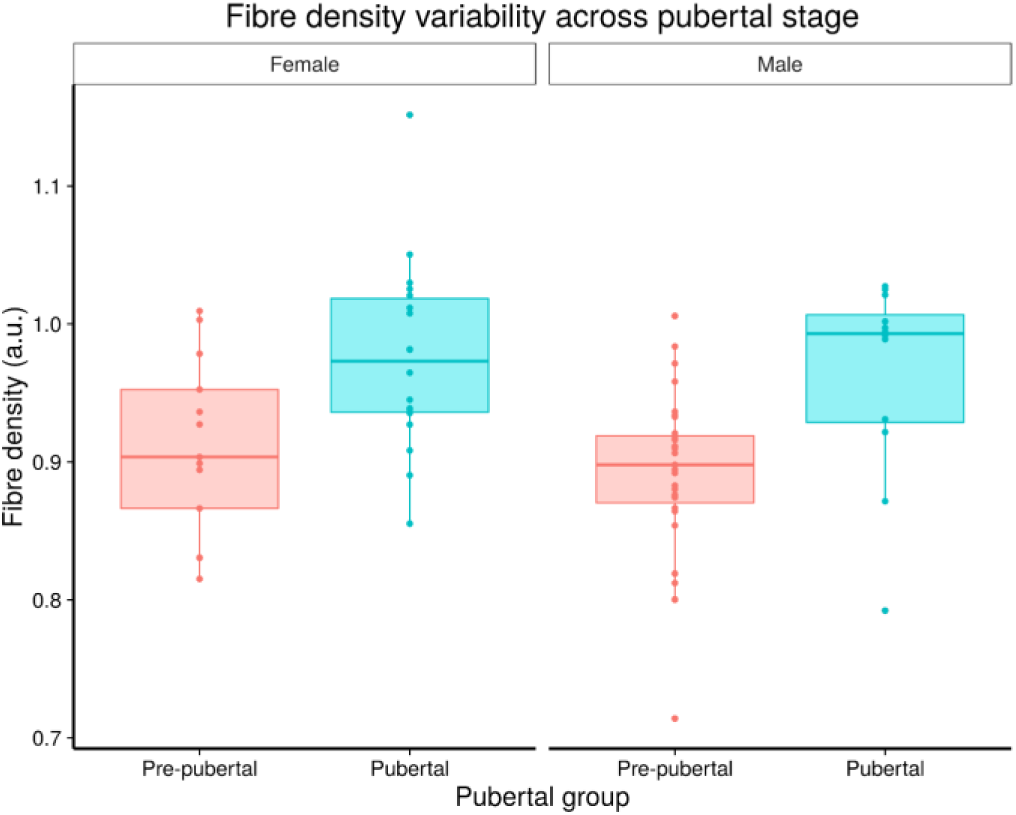
Spread of fibre density in each pubertal group faceted by sex.

